# Identification of Fusarium Head Blight resistance in *Triticum timopheevii* accessions and characterisation of wheat-*T. timopheevii* introgression lines for enhanced resistance

**DOI:** 10.1101/2022.05.13.491811

**Authors:** Andrew Steed, Julie King, Surbhi Grewal, Cai-yun Yang, Martha Clarke, Ian King, Paul Nicholson

## Abstract

A diverse panel of wheat wild relative species was screened for resistance to Fusarium head blight (FHB) by spray inoculation. The great majority of species and accessions were susceptible or highly susceptible to FHB. Accessions of *Triticum timopheevii* (P95-99.1-1), *Agropyron desertorum* (9439957) and *Elymus vaillantianus* (531552) were highly resistant to FHB while additional accessions of *T. timopheevii* were found to be susceptible to FHB. A combination of spray and point inoculation assessments over two consecutive seasons indicated that the resistance in accession P95-99.1-1 was due to enhanced resistance to initial infection of the fungus (type 1 resistance), and not to reduction in spread (type 2 resistance).

A panel of wheat-*T. timopheevii* (accession P95-99.1-1) introgression lines was screened for FHB resistance over two consecutive seasons using spray inoculation. Most introgression lines were similar in susceptibility to FHB as the wheat recipient (Paragon) but substitution of the terminal portion of chromosome 3BS of wheat with a similar-sized portion of 3G of *T. timopheevii* significantly enhanced FHB resistance in the wheat background.

## Introduction

Fusarium head blight (FHB) is a highly damaging disease of bread wheat (*Triticum aestivum*) and durum wheat *(T. durum*) in many parts of the world. Infection generally occurs during flowering when susceptibility to FHB is greatest in the host (Franco et al., 2021) leading to yield loss and reduced grain quality (Spanic et al., 2021). The disease is mainly caused by *Fusarium graminearum* sensu stricto but other species including *F. culmorum* and *F. asiaticum* can be important in some regions (Valverde-Bogantes et al., 2020). These species can produce trichothecene mycotoxins such as deoxynivalenol (DON) and nivalenol (NIV) that contaminate grain and pose a risk to human and animal consumers (Amarasinghe et al., 2019). It is widely accepted that the use of FHB-resistant varieties is the most effective and sustainable means to mitigate against losses caused by FHB (Bai et al., 2018). Breeding for resistance to FHB is particularly challenging because of the generally polygenic nature of resistance, high level of genotype-by-environment interactions and the high cost of phenotyping.

Resistance to FHB was originally differentiated into two classes: resistance to initial infection (type 1) and resistance to spread within the spike (type 2) (Schroeder and Christensen, 1963). Additional classes of resistance have been proposed including degradation of DON (Miller et al., 1985) and DON tolerance being grouped and termed type 3 resistance (Mesterhazy et al., 1999) and resistance to kernel infection as type 4 (Mesterhazy et al., 1999). Type 1 resistance is determined by spraying spikes at mid-anthesis with a conidial suspension and measuring the percentage of diseased spikes whereas type 2 resistance is determined by inoculating single florets with conidial inoculum and measuring the number or percentage of diseased spikelets over time.

Resistance to FHB is quantitatively inherited and over 100 quantitative trait loci (QTL) have been reported to date distributed across all 21 chromosomes of bread wheat (Buerstmayr et al., 2020). Many potent QTL have been identified in Asian germplasm including Sumai 3, Wangshuibai, Nobeokabouzu and Nyu Bai (Buerstmayr et al., 2020). The most highly studied source of resistance is the Chinese variety Sumai 3 in which three QTL were originally identified on the short arms of chromosomes 3B and 5A and the long arm of chromosome 6B (Buerstmayr et al., 2020).

Most studies have focussed on assessing type 2 resistance because it is more stable than type 1 resistance and less prone to influence by environmental factors. Simple Mendelian inheritance has been demonstrated for a number of QTL when isolated into susceptible wheat variety backgrounds (Cuthbert et al., 2006; Cuthbert et al., 2007). Seven such QTL have been formally recognised as genes *Fhb1*-*Fhb7* (Guo et al., 2015). The identity of *Fhb1* (type 2 resistance) was originally identified as a chimeric lectin with agglutinin domains and a pore forming toxin domain (Rawat et al., 2016) but later studies have cast doubt on this with a second gene being proposed to be responsible. A histidine-rich calcium-binding protein was demonstrated to provide resistance to FHB by two groups although one group (Su et al., 2019) concluded that the resistance is due to the loss of function while the second group (Li et al., 2019) concluded that resistance was the result of a gain of function.

The search for additional sources of potent FHB resistance continues and extends beyond the primary gene-pool into wheat relatives. Resistance has been identified in a number of chromosome segments introgressed into wheat from wild relatives. In some instances, these resistances have been considered as genes due to the lack, or extremely limited degree, of recombination between the introgressed segment and wheat chromosome. Introgression of a portion of chromosome 3St of *Elymus repens* into 3D of wheat confers high levels of type 2 resistance (Fedak et al., 2017; Gong et al., 2019). Substitution of the short arm of chromosome 7AS of wheat with the short arm of *Leymus racemosus* 7Lr#1 provides a high level of type 2 resistance and this has been designated as *Fhb3* (Qi et al., 2008). Similarly, replacement of the short arm of chromosome 1A (1AS) of wheat with the short arm of chromosome 1E^ts^#1S of *Elymus tsukushiensis* also significantly enhances type 2 resistance in wheat and has been designated as *Fhb6* (Cainong et al., 2015). The most extensively studied resistance in a wheat relative derives from *Thinopyrum elongatum* (*Th. ponticum*) (Guo et al., 2015; Ceoloni et al., 2017). Substitution of the long arm of wheat chromosome 7D (7DL) with the long arm of 7El_2_ of *Th. elongatum* confers very high levels of type 2 resistance (Shen and Ohm, 2007; Zhang et al., 2011; Wang et al., 2020). The gene responsible for this resistance, termed *Fhb7* (formerly *Fhblop*), has recently been isolated and shown to encode a glutathione S-transferase (GST) that functions through de-epoxidation of trichothecenes (Wang et al., 2020).

The objectives of the present study were to: 1) screen accessions of wheat relatives to identify FHB resistance, 2) determine whether resistance is predominantly of type 1 (resistance to initial infection) or type 2 (resistance to spread in the spike), 3) determine whether segments of chromosomes from resistant wheat relatives conferred FHB resistance when introgressed into wheat.

## Methods

### Fungal materials

All Fusarium isolates used in this study originated from the UK and are kept as part of the JIC facultative pathogen collection. Isolates were maintained as reported previously (Hales et al., 2020).

### Wheat wild relative species FHB screen

A diverse panel of wheat wild relative accessions from Nottingham/BBSRC Wheat Research Centre (originally obtained from the Germplasm Resource Unit (GRU) at the JIC and the United States Department of Agriculture (USDA)) was screened for FHB resistance. Material was sown in the winter of 2011 and given natural vernalization in an unheated, unlit glasshouse. In the spring of 2012 seedlings were transplanted into 1 litre pots of cereals mix and grown in a Keder Greenhouse until mid-anthesis. Due to the diverse growth habits and morphology of the material it was spray inoculated repeatedly around the time of mid-anthesis to ensure that all the material received inoculum at the period of maximum susceptibility. The inoculum consisted of conidia (1×10^5^ conidia ml^-1^) of a DON producing isolate of *F. culmorum*, applied using a handheld mister.

The spikes of some species contain very few spikelets making it inappropriate to use a conventional scoring system based upon the percentage of spikelets with symptoms. Disease levels were assessed between three and four weeks post inoculation using a 1-9 rating based upon a combination of percentage of spikes affected and percentage of spikelets showing disease in infected spikes. A visual disease score of 1 indicating “no visible disease”, a visual disease score of 9 indicating very high disease levels of 90-100 % infected spikelets on all spikes at or near mid-anthesis at the time of inoculation.

### FHB disease assessment of *T. timopheevii* accessions by spray inoculation (2015 and 2016)

Seven *T. timopheevii* accessions (all obtained from the USDA) and a susceptible wheat variety (Highbury – obtained from the GRU) were assessed for FHB resistance in 2015 by spray inoculation with conidia (1×10^5^ ml^-1^) of a DON producing isolate of *F. graminearum*. Between 11 and 39 individual spikes per line from multiple plants were spray inoculated at mid-anthesis, and disease was assessed at 21 days post inoculation (dpi) and the percentage of infected spikelets per spike calculated. All statistical analysis used Genstat 18^th^. The trial was unblocked, and GLM analysis used “Inoculation date” and “Line” in the model. GLM was used to calculate predicted means and standard errors for percentage of infected spikelets for each line.

Three *T. timopheevii* accessions, a wheat/Tim_P95-99.1-1 amphidiploid and the wheat variety Highbury (parent to the amphidiploid) were assessed for FHB resistance in 2016 by spray inoculation with conidia (1×10^5^ ml^-1^) of a DON producing isolate of *F. graminearum*. The trial had a randomised block design containing six replicate blocks with 5-8 plants per line. Multiple ears per plant were spray inoculated at mid-anthesis, and disease was assessed at 19dpi and the percentage of infected spikelets per spike calculated. All statistical analysis used Genstat 18^th^, GLM analysis used “Inoculation date”, “Replicate” and “Line” in the model. Using GLM, predicted means and standard errors were calculated or each line.

### FHB disease assessment of *T. timopheevii* accessions by point inoculation (2015 and 2016)

Seven *T. timopheevii* accessions and a susceptible wheat variety (Highbury) were assessed for FHB resistance following point inoculation in 2015. Between 10 and 27 individual spikes per line from multiple plants were point inoculated at mid-anthesis. Inoculum (10µl) of a DON producing *F. graminearum* isolate (1×10^6^ conidia ml^-1^) was introduced directly into a central spikelet for each spike. Disease was assessed at 21dpi and the number of infected spikelets above and below the point of inoculation recorded. All statistical analysis used Genstat 18^th^. The trial was unblocked, and GLM analysis used “Inoculation date” and “Line” in the model. GLM was used to calculate predicted means and standard errors for the number of infected spikelets above and below the point of inoculation for each line.

Three *T. timopheevii* accessions, a wheat/Tim_P95-99.1-1 amphidiploid and the wheat variety Highbury (parent to the amphidiploid) were assessed for FHB resistance following point inoculation in 2016. The trial had a randomised block design containing 6 replicate blocks with 5-8 plants for each line with multiple spikes inoculated for each plant. Individual spikes were point inoculated into the central floret at mid-anthesis with *F. graminearum* at 1×10^−6^ spore per ml, using a 0.5 ml insulin syringe. Disease was assessed at 14dpi and the number of infected spikelets above and below the point of inoculation recorded. All statistical analysis used Genstat 18^th^, GLM analysis used “Inoculation date” “Replicate” and “Line” in the model. GLM was used to calculate predicted means and standard errors for the number of infected spikelets above and below the point of inoculation for each line.

### Development of wheat/*T. timopheevii* (P95-99.1-1) introgression lines

Development of wheat/*T. timopheevii* introgression lines was as outlined in Devi et al. (2019) and King et al (2022). Briefly, Paragon *ph1/ph1* (obtained from the GRU) was used as the female parent in a cross with *T. timopheevii* P95-99.1-1 to generate F_1_ interspecific hybrids. The F_1_ hybrids were then backcrossed to Paragon to generate BC_1_, BC_2_, BC_3_ and BC_4_ plants. Molecular characterisation of the introgression lines was initially carried out using the Axiom^®^ Wheat-Relative Genotyping Array (Devi et al., 2019). When genotyping of these plants showed the number of introgressions present to be three or less, the plants were then self-fertilised. Chromosome-specific KASP markers, polymorphic between wheat and *T. timopheevii* have been developed at the WRC and 480 of these KASP markers have been used to characterise a panel of homozygous wheat/*T. timopheevii* introgression lines including those investigated in this work (King et al. 2022).

### Fluorescence *in situ* hybridisation (FISH)

FISH analysis of wheat-*T. timopheevii* introgression lines was carried out as described in (2022 ref). Root metaphase spreads of chromosomes were hybridised with probes pSc119.2 (McIntyre et al., 1990) and pAs.1 (Rayburn and Gill, 1986) that were nick-labelled (Rigby et al., 1977) with Alexa Fluor 488-5-dUTP (green) and Alexa Fluor 594-5-dUTP (red), respectively. Karyotyping of labelled chromosomes was done in accordance with the nomenclature reported by Badaeva et al. 2016.

### FHB disease assessment of wheat-*T. timopheevii* introgression lines by spray inoculation (2020 and 2021)

Twenty-five wheat-*T. timopheevii* introgression lines and wheat variety Paragon were assessed for FHB resistance by spray inoculation in a Keder Greenhouse in 2020 as described above. The trial had a randomised block design with between 4 and 6 individual plants distributed within 3 replicate blocks. Multiple spikes per plant were spray inoculated at mid-anthesis with conidia (1×10^5^ ml^-1^) of a DON producing isolate of *F. culmorum* using a handheld mister.

Twenty-nine wheat-*T. timopheevii* introgression lines and FHB susceptible wheat varieties Highbury and Paragon were assessed for FHB resistance by spray inoculation in a Keder Greenhouse in 2021 as described above. The trial had a randomised block design with between 5 and 10 individual plants distributed within 4 replicate blocks. Spikes were inoculated at mid-anthesis as described above.

Disease in both trials was assessed at 21dpi and the percentage of infected spikelets per ear calculated. All statistical analysis used Genstat 18^th^, GLM analysis used “Inoculation date”, “Replicate” and “Line” in the model. Using GLM, predicted means and standard errors were calculated for each line.

## Results

### FHB screen of wheat relatives

The initial screen of 113 wheat wild relative accessions revealed that most accessions of all species were susceptible or highly susceptible to FHB (Table 1). Where more than one accession of a species was tested most showed similar levels of FHB susceptibility but evidence of variation in FHB susceptibility within species was observed for some species. One accession of *Agropyron desertorum* (PI 439957) was highly resistant to FHB (FHB score 2) while a second accession (PI 439953) was highly susceptible (FHB score 8). Similarly, accession 2060002 of *Aegilops biuncialis* was moderately resistant (FHB score 3) while accession 2060003 was highly susceptible (FHB score 9). The majority of accessions of *Aegilops sharonensis* were highly susceptible to FHB (FHB score 8-9) but a few (AS_01512//8, AS_01850//17 and AS_01930//19) were moderately resistant (FHB score 4-5). The single accession of *Elymus vaillantianus* (PI 531552) was highly resistant to FHB (FHB score 2) and exhibited few symptoms. The single accession of *Triticum timopheevii* (Tim_P95-99.1-1) was notable as it exhibited a very high level of resistance to FHB (Table 1) with no symptoms being apparent even at later dates post inoculation.

**Table 1.**
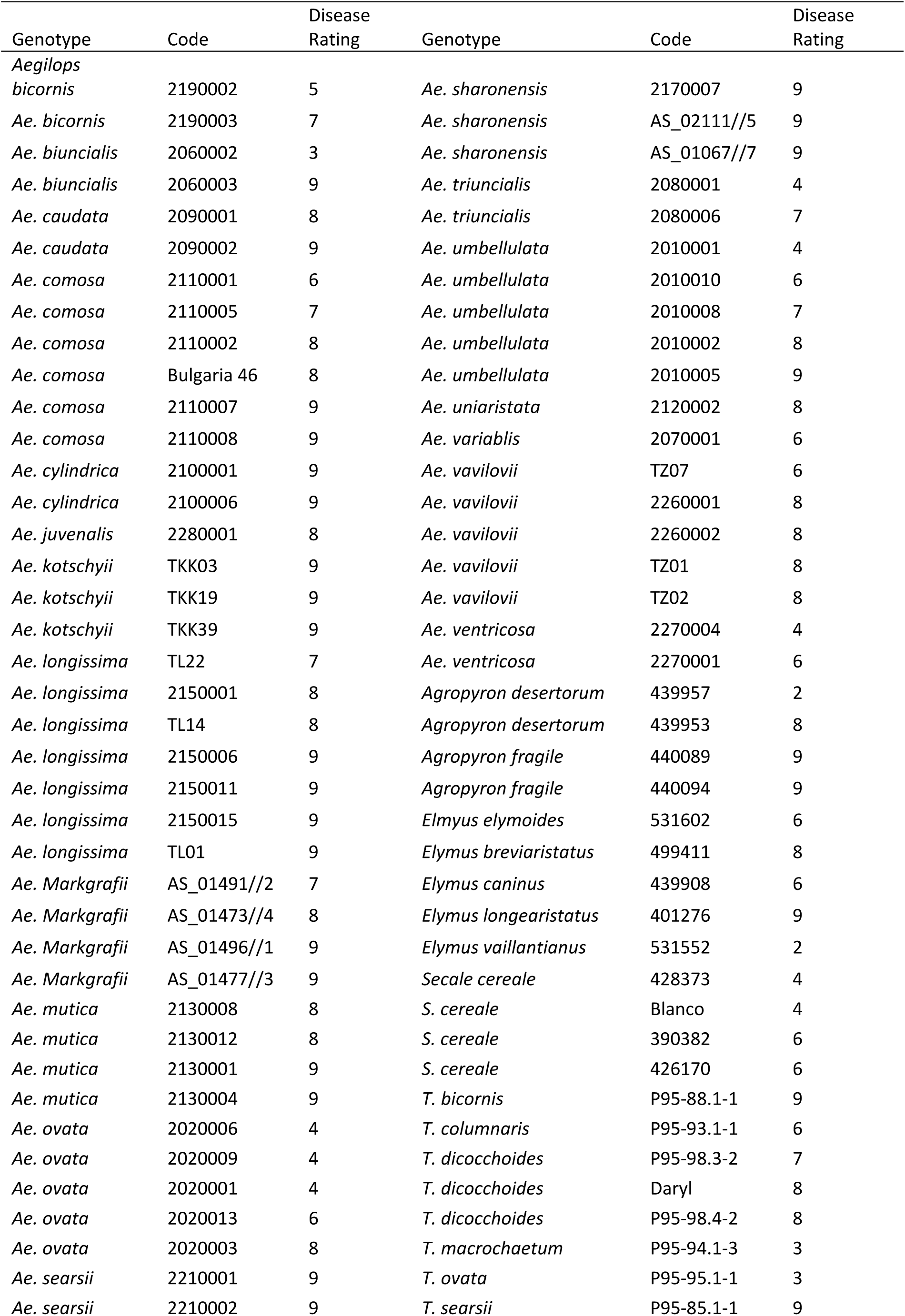
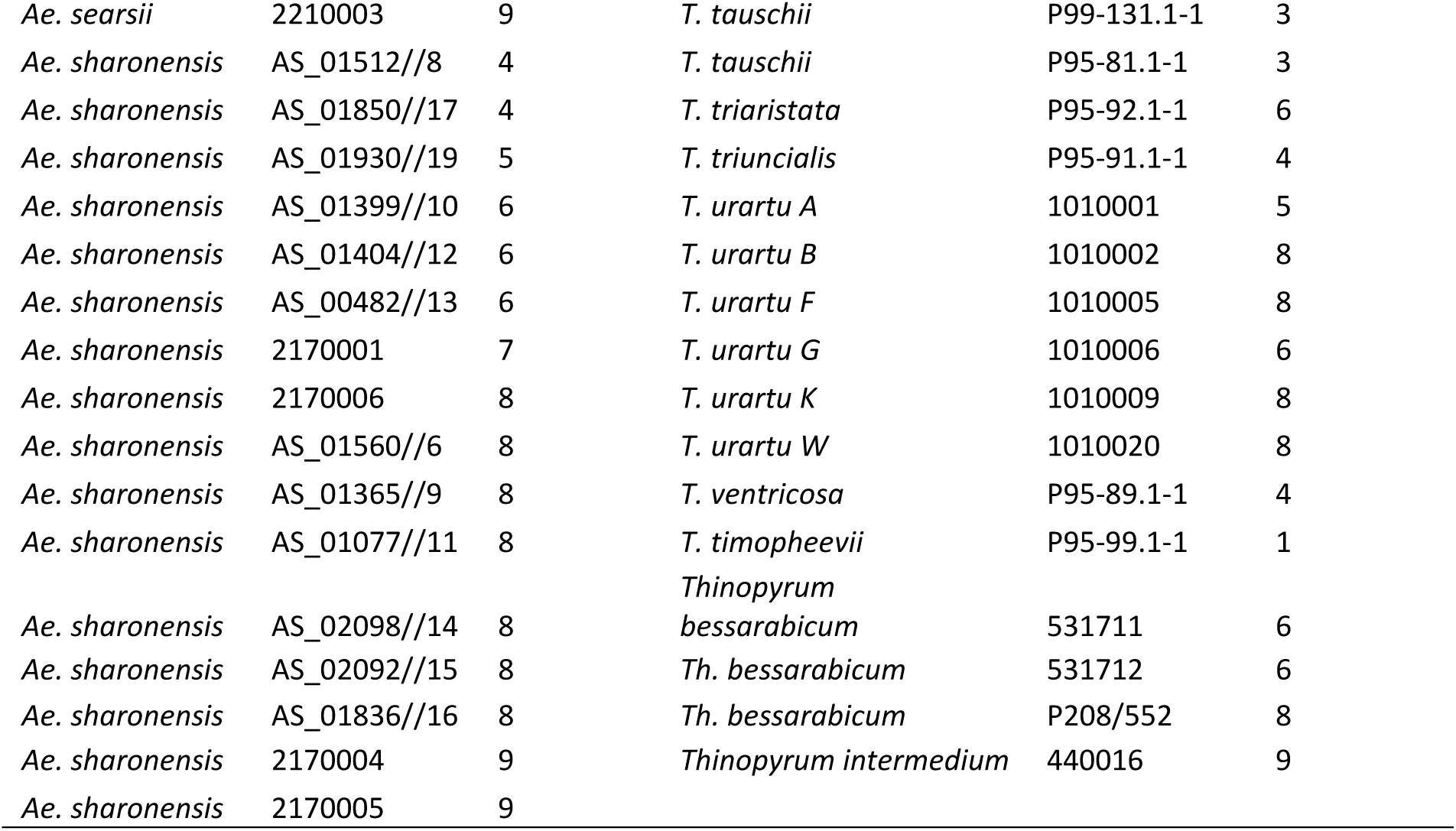
Fusarium head blight disease rating of 113 wild grass species/accessions of wheat relatives following spray inoculation with conidia of *F. graminearum*

### FHB spray inoculation screening of accessions of *T. timopheevii*

Given the variation in FHB resistance observed in many species in the initial screen, six additional accessions of *T. timopheevii* were obtained to compare their resistance with that of accession Tim_P95-99.1-1. Resistance to FHB is highly sensitive to environmental factors so Tim_P95-99.1-1 and selected additional accessions were tested across two seasons. Resistance derived from wheat relatives may not always be effective when introgressed into wheat (Innes and Kerber, 1994; Rines et al., 2007). An amphidiploid line was produced by crossing Tim_P95-99.1-1 to the FHB-susceptible wheat variety Highbury. The amphidiploid line was only available for FHB screening in 2016.

FHB spray inoculation of six additional *T. timopheevii* accessions alongside Tim_P95-99.1-1 and the susceptible wheat variety Highbury was undertaken in 2015. At 21 dpi, over 84% of spikelets of Highbury were symptomatic for FHB. All accessions of *T. timopheevii*, with the exception of Tim_427998, were significantly more resistant (P<0.001) than Highbury. Tim_P95-99.1-1 exhibited the greatest level of FHB resistance (29% spikelets infected) with a lower disease score than all the other *T. timopheevii* accessions tested. Indeed, the disease score for Tim_P95-99.1-1 was significantly lower (P< .001) than four of the *T. timopheevii* accessions and the wheat variety Highbury (Figure 1). Two other *T. timopheevii* accessions (Tim_PI_289752 and Tim_PI_427414) were also markedly more resistant than Highbury with only 45% and 45.8% of spikelets infected respectively.

**Figure 1.**
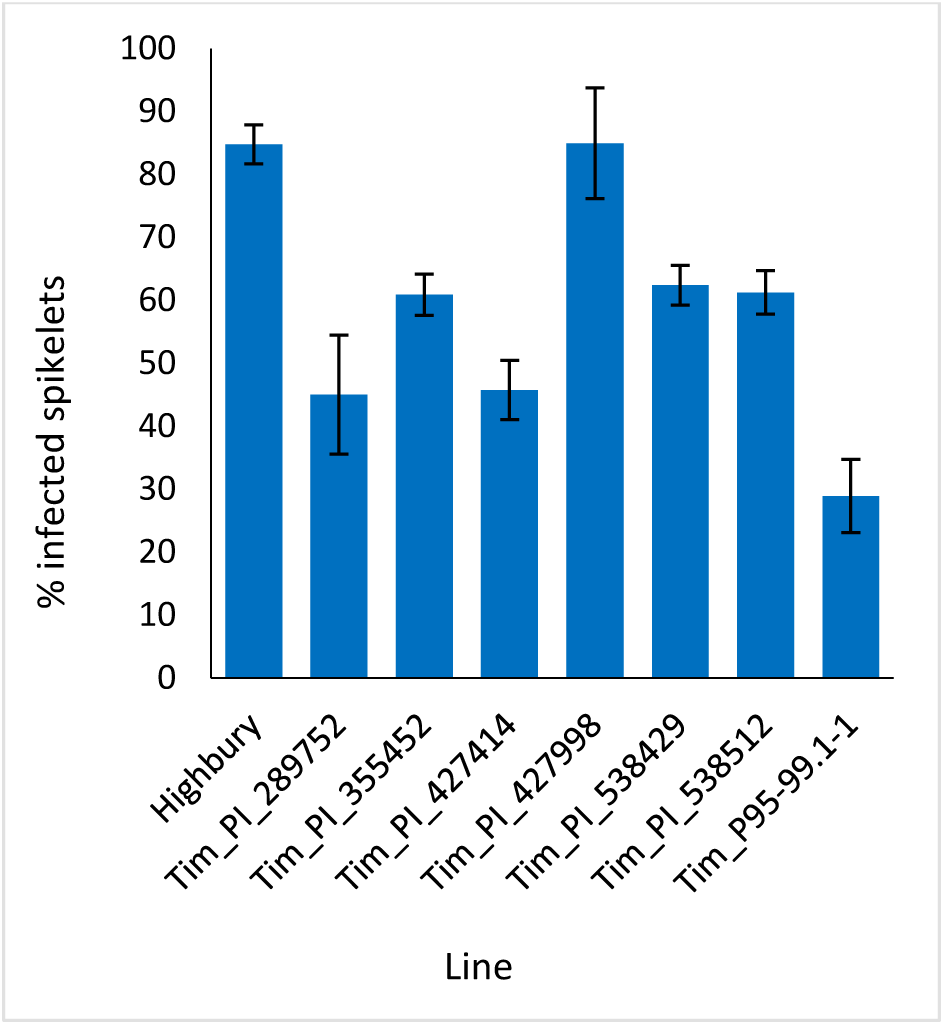
Visual FHB disease score from 2015 trial of seven *T. timopheevii* accessions and Highbury susceptible control expressed as percentage of infected spikelets 21 days post spray inoculation with *F. graminearum* (1×10^5^ conidia per ml).

In the 2016 FHB spray screen disease levels were not as high as in 2015. Highbury had approximately 45% of spikelets exhibiting disease. In contrast, Tim_P95-99.1-1 had an extremely low level of disease with only 1.3% of spikelets exhibiting symptoms (Figure 2). As in 2015, accessions Tim_PI_538429 and Tim_PI_538512 were significantly less diseased (P<0.001) than Highbury with 18.8% and 11% of spikelets infected. Disease levels in the amphidiploid were also very low with an average of 5% of spikelets infected and were not statistically different (P=0.41) from those in the Tim_95-99.1-1 parent. These results reveal that the FHB resistance in Tim-P95-99.1-1 is stable and is expressed in a wheat background.

**Figure 2.**
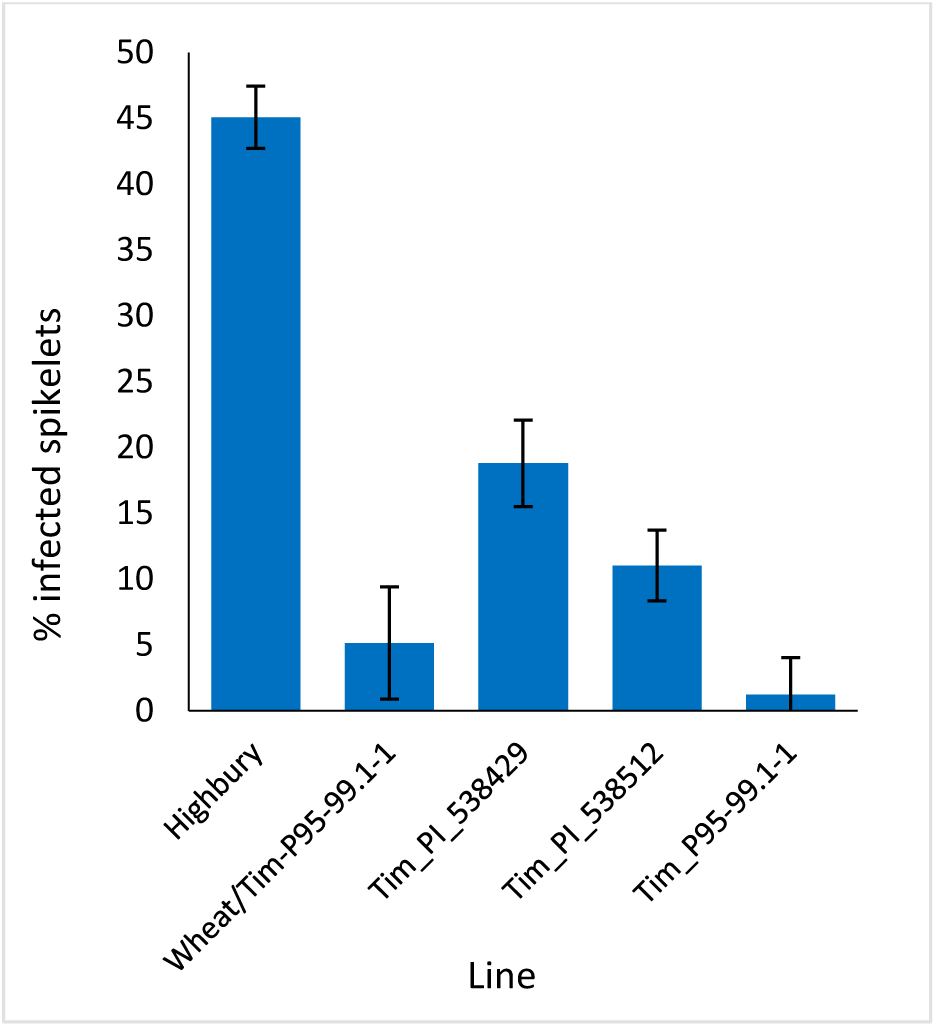
Visual FHB disease score from 2016 trial of three *T. timopheevii* accessions, *T. timopheevii* x Highbury amphidiploid and wheat variety Highbury expressed as percentage of infected spikelets 14 days post spray inoculation with *F. graminearum* (1×10^5^ conidia per ml).

### FHB point inoculation screening of accessions of *T. timopheevii*

The preliminary FHB screen involved spray inoculation which reveals overall levels of FHB resistance. Point inoculation is used to determine whether the resistance is of type 2, resistance to spread. Tim_P95-99.1-1 along with six other *T. timopheevii* accessions and the wheat variety Highbury were point inoculated at mid-anthesis. Symptoms above the point of inoculation generally reflect susceptibility to the effects of DON while those below reflect colonisation by the pathogen. For these reasons, disease above and below the point of inoculation was assessed separately. At 21 dpi disease levels in Highbury were 18% and 32.7% infected spikelets above and below the point of inoculation respectively (Figure 3). Four accessions of *T. timopheevii* (Tim_PI_289752, Tim_PI_427414, Tim_PI_ 538429, Tim_PI_538512) were significantly more susceptible to FHB than Highbury for symptoms both below and above the point of inoculation. None of the *T. timopheevii* accessions exhibited greater resistance to spread of symptoms either above or below the inoculation point than Highbury. Tim_538429 was extremely susceptible to spread of symptoms both above and below the inoculation point indicating an inability to restrict fungal colonisation and a high level of susceptibility to the effects of DON (Figure 3). Accessions Tim_PI_289752 and Tim_PI_427414 exhibited high disease levels below the point of inoculation indicating that they lack the ability to restrict fungal colonisation. Disease above and below the point of inoculation in the other accessions, including Tim_P95.99.1-1, were slightly, but not significantly, more susceptible than the wheat variety Highbury. Overall, no evidence was apparent that indicated that any of the *T. timopheevii* accessions possessed greater levels of type 2 resistance than Highbury.

**Figure 3.**
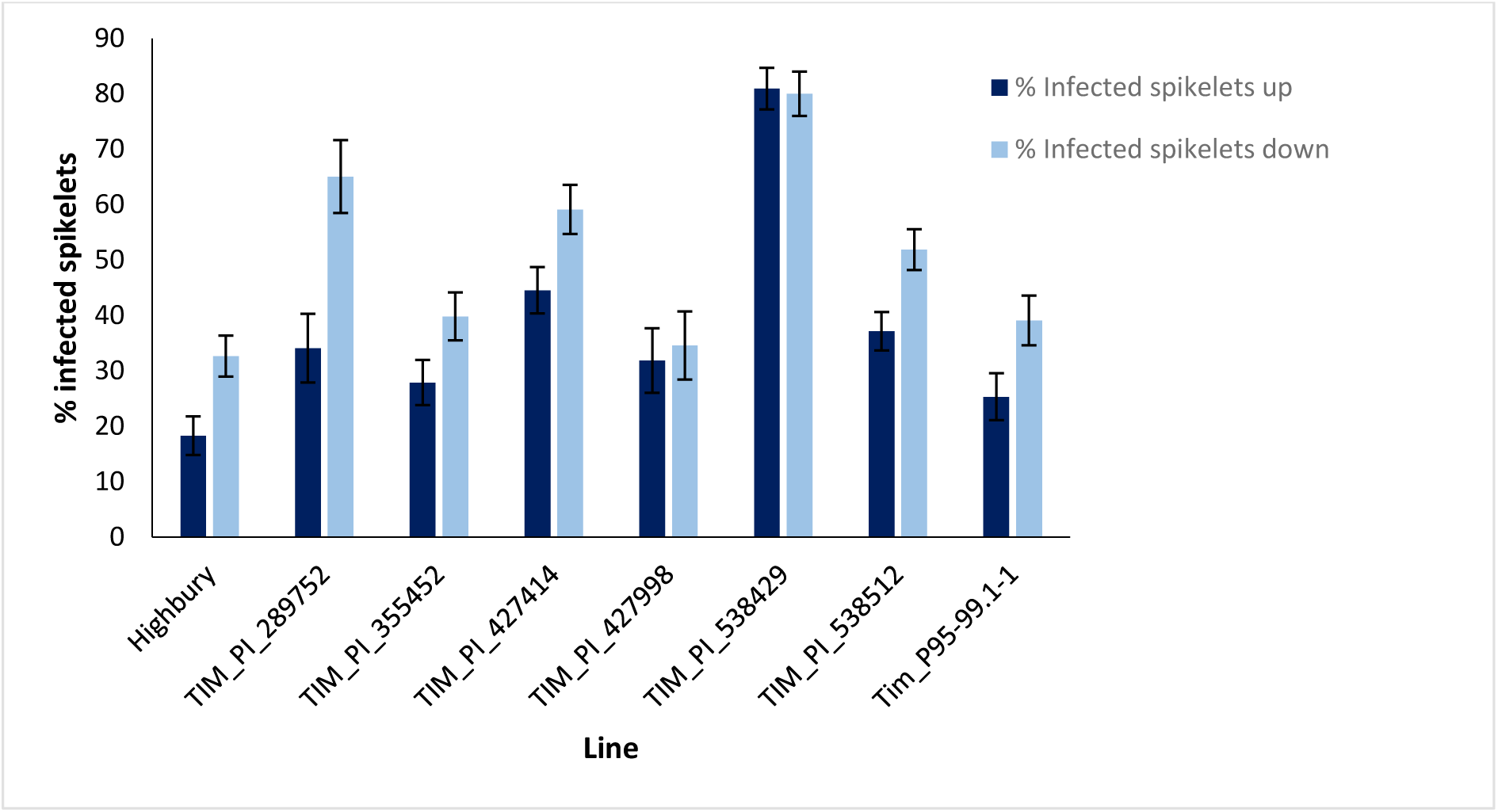
Visual FHB disease score from 2015 trial of seven *T. timopheevii* accessions and Highbury susceptible control expressed as number of infected spikelets above and below the point of inoculation, 21 days post inoculation with *F. graminearum* (1×10^6^ conidia per ml).

Disease progress in the point inoculation screen in 2016 was greater than in the previous year with disease levels in Highbury of 75.5% and 68.2% above and below the inoculation point respectively (Figure 4). As in the 2015 screen, disease symptoms above and below the inoculation point in Tim_PI_538429 were significantly greater (P=0.028 and P=0.05 respectively) than those in Highbury although the differential was markedly less than that in the earlier screen where disease pressure was lower (compare Figure 3 and Figure 4). Unlike in the earlier trial, disease levels above and below the inoculation point in Tim_PI_538512 were not significantly greater than those in Highbury with disease level above the point of inoculation being significantly less (P<0.001) than that of Highbury (Figure 4).

**Figure 4.**
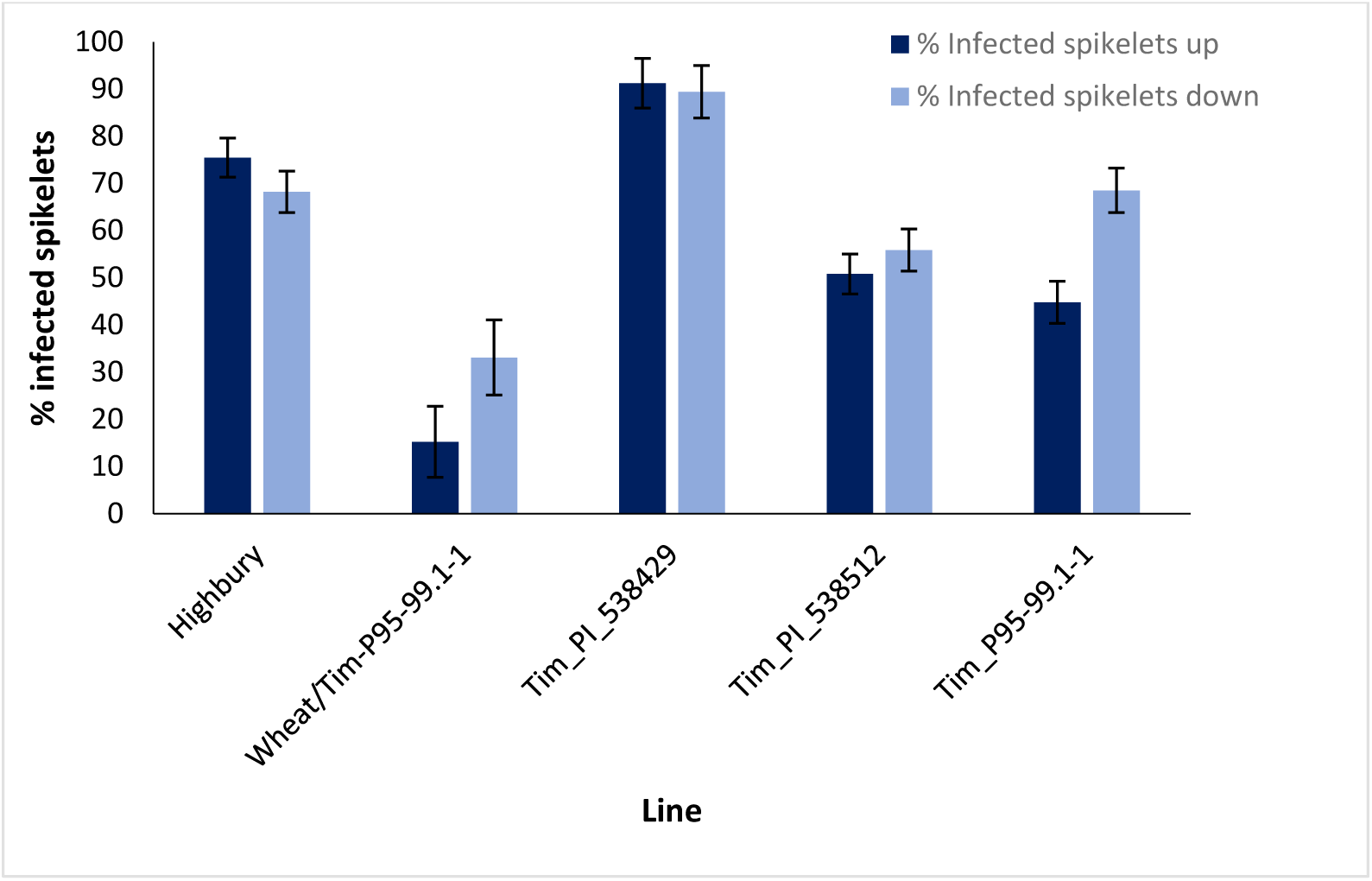
Visual FHB disease score from 2016 trial of three *T. timopheevii* accessions, *T. timopheevii* x Highbury amphidiploid and wheat variety Highbury expressed as number of infected spikelets above and below the point of inoculation, 14 days post inoculation with *F. graminearum* (1×10^6^ conidia per ml).

Symptoms below the inoculation point in Tim_P95-99.1-1 were similar to those in Highbury (P=0.96) while bleaching symptoms above the point of inoculation were significantly less than those in Highbury (P<0.001). Despite this, the differential in disease levels between Tim_P95-99.1-1 and Highbury were very much less following point inoculation than after spray inoculation (compare Figure 2 and Figure 4). Unexpectedly, disease levels above (15.2%) and below (33.1%) the point of inoculation were significantly less (P<0.001) in the amphidiploid line than either parent. It was noted that some spikes of the amphidiploid were partially sterile, and this may have reduced susceptibility in this line.

### FHB screening of wheat lines carrying introgressions from Tim_P95-99.1-1

The above studies revealed that accession Tim_P95-99.1-1 possesses a very high level of resistance to FHB and that this resistance is predominantly of type 1 rather than type 2. The studies also demonstrated that the FHB resistance in Tim_P95-99.1-1 was expressed in an amphidiploid line produced by crossing to the wheat variety Highbury. We next investigated whether introgression of Tim_P95-99.1-1 chromosome segments into wheat would confer any level of increased resistance to FHB. The development of a panel of introgressions of Tim_P95-99.1-1 into the wheat variety Paragon has been reported previously (Devi et al., 2019). These lines were advanced and a selection of 32 homozygous introgression lines from this panel (King et al., 2022; Table 2) were screened for FHB resistance by spray inoculation. Twenty-five of these introgression lines were tested in 2020 and 29 lines in 2021. Each line was subjected to genotyping using chromosome-specific KASP markers (King et al., 2022). These introgression lines contain a variety of segments from each of the two subgenomes present in *T. timopheevii* (A^t^ and G) (Fig 7; Table 2).

**Table 2.** Chromosomal segments of *T. timopheevii* in Paragon wheat introgression lines

| Line | segment |  |  |  |  | comments |
| --- | --- | --- | --- | --- | --- | --- |
|  | 1 | 2 | 3 | 4 | 5 |  |
| BC2F4-40 | 3G | - | - | - | - |  |
| Tim1 | 2A<sup>t</sup> | 6A<sup>t</sup> | 7G | - | - |  |
| Tim2 | 1A<sup>t</sup> | 3G | 7G | - | - | lacks 3BS |
| Tim3 | 2A<sup>t</sup> | 5A<sup>t</sup> | 7A<sup>t</sup> | - | - |  |
| Tim4 | 5A<sup>t*</sup> | 6G | 7G | - | - |  |
| Tim5 | 2G | 3G | - | - | - |  |
| Tim6 | 2G | 3G | 6G | - | - |  |
| Tim7 | 2A<sup>t</sup> | 7G | - | - | - |  |
| Tim8 | 2A<sup>t</sup> | 5G | - | - | - |  |
| Tim9 | 2A<sup>t</sup> | 5G | 7A<sup>t</sup> | 7G | - |  |
| Tim10 | 2A<sup>t</sup> | 2G | 3A<sup>t</sup> | 5At | 5G |  |
| Tim11 | 2G | 6A<sup>t</sup> | - | - | - |  |
| Tim12 | 2G | 6A<sup>t</sup> | 7G | - | - |  |
| Tim13 | 7A<sup>t*</sup> | 7G | - | - | - |  |
| Tim14 | 1A<sup>t</sup> | 2A<sup>t</sup> | 4G | - | - |  |
| Tim15 | 1A<sup>t</sup> | 1G | - | - | - |  |
| Tim17 | 4G | 5A<sup>t</sup> | - | - | - |  |
| Tim18 | 5G | 6A<sup>t</sup> | 6G | - | - |  |
| Tim19 | 1G | 4G | 6A<sup>t</sup> | - | - |  |
| Tim21 | 2A<sup>t</sup> | 4G | - | - | - |  |
| Tim22 | 6A<sup>t</sup> | - | - | - | - |  |
| Tim23 | 5A<sup>t</sup> | - | - | - | - |  |
| Tim24 | 3G | 7G | - | - | - |  |
| Tim25 | 2G | 5A<sup>t</sup> | - | - | - |  |
| Tim26 | 2G | - | - | - | - |  |
| Tim27 | 2G | 3G | 5A<sup>t</sup> | 6G | - |  |
| Tim28 | 1A<sup>t</sup> | 5G | 6A<sup>t</sup> | - | - |  |
| Tim29 | 1A<sup>t</sup> | 2A<sup>t</sup> | 4G | - | - |  |
| Tim30 | 2A<sup>t</sup> | - | - | - | - |  |
| Tim31 | 5A<sup>t</sup> | 6G | - | - | - |  |
| Tim33 | 2G | 7A<sup>t</sup> | - | - | - |  |
| Tim35 | 1A<sup>t</sup> | 3G | 7G | - | - |  |

\* two segments

In 2020, disease levels on Paragon wheat were moderate with 47.5% spikelets exhibiting FHB symptoms (Figure 5). Significant differences were observed in the levels of FHB resistance among the introgression lines. Two introgression lines (Tim7 and Tim12) exhibited significantly higher disease levels than the Paragon recipient, 73.5% and 88.4% respectively. Two introgression lines, Tim6 and Tim5 appeared highly resistant to FHB with disease levels of 4% and 10% respectively and were significantly more resistant than Paragon (P<0.001) (Fig 5). These two lines were of interest as although they both contained introgressed segments from linkage groups 2G and 3G of *T. timopheevii*, they were unique in the set tested as containing segments from 3G (Figure 7). The presence of segments of 3G was confirmed with fluorescent *in situ* hybridisation (FISH; Figure 8). Three other introgression lines (Tim26, Tim28 and Tim11) were also significantly more resistant than Paragon, but to a lesser extent, with 22.9%, 23.2% and 26.9% spikelets infected respectively.

**Figure 5.**
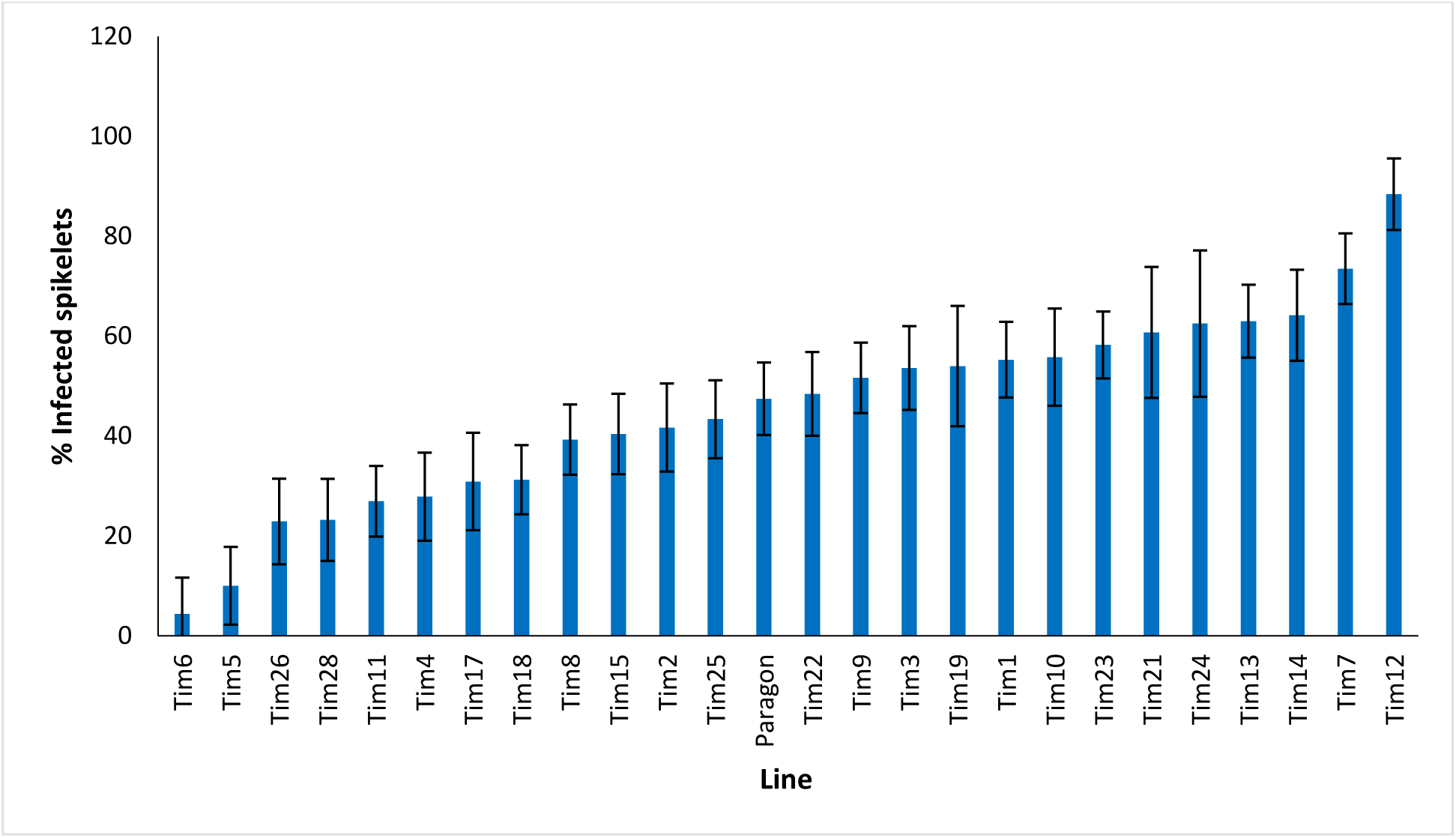
Visual FHB disease score from 2020 trial of 25 *T. timopheevii* accession P95-99.1-1/wheat introgression lines and Paragon susceptible control, expressed as percentage of infected spikelets 21 days post spray inoculation with *F. culmorum* (1×10^5^ conidia per ml).

Disease levels were higher in the 2021 screen with 58.6% of spikelets of Paragon exhibiting FHB symptoms. No introgression line appeared more susceptible to FHB than Paragon. Four introgression lines Tim11, (22.4%) BC_2_F_4_-40 (23.6%), Tim33 (24.6%) and Tim5 (25.5%) were significantly (P<0.001) more resistant to FHB than Paragon. Both Tim11 and Tim5 had also shown significantly greater resistance to FHB in the screen in the previous year. Neither BC_2_F_4_-40 or Tim33 had been included in the screen in the previous year. Two additional introgression lines (Tim29 and Tim30) also exhibited significantly less disease than Paragon but significantly more than the above-mentioned four lines.

## Discussion

Several sources of potent FHB resistance have been identified in *Triticum aestivum*, particularly in lines from China and Japan (Buerstmayr et al., 2009). Subsequent study revealed that, in many cases, the genetic basis of resistance was similar and so research continues to expand the range of FHB resistance available to wheat breeders. Potentially useful resistance to FHB has been identified in both the secondary and tertiary gene pools (Steed et al 2005; Ceoloni et al 2017). In the present work, accessions of three species exhibited very high levels of FHB resistance. While resistance in *Agropyron desertorum* appears not to have been reported previously FHB resistance has been identified in both *E. vaillantianus* (syn. *E repens*) (Fedak et al., 2017; Gong et al., 2019) and *T. timopheevii* (Malihipour et al., 2016). In the present study only one accession of *T. timopheevii* (P95-99.1-1) exhibited resistance to FHB while the other accessions were moderately to highly susceptible. *Triticum timopheevii* is an allopolyploid (2n=4x=28) comprising the A^t^ genome similar to that of the A genome progenitor of wheat (*Triticum aestivum*) and the G genome being more similar to the B genome of *T. aestivum* (Dvorak and Zhang, 1990: Dvorak et al., 1993; Devi, 2019). Recombination between the A and A^t^ genomes is more frequent than that between the B and G genomes reflecting their relative relatedness (Feldman, 1966; Devi et al., 2019).

Resistance identified in *T. timopheevii* to several pathogens has been introduced into wheat. Genes for resistance to stem rust include *Sr36, Sr40*, and *Sr50* on chromosome 2B (Brown-Guedira et al., 2003) and *Sr37* on 4B (Bai et al., 1998). Resistance to leaf rust has been introduced on 5B (*Lr18*) (Sadeghabad et al., 2017)and 5A (un-named resistance) (Bai et al., 1998). Resistance conditioned by both leaf rust genes was recessive in bread wheat while the 5A resistance was dominant in durum wheat indicating that the expression of resistance is dependent upon the background (Bai et al., 1998). Resistance to powdery mildew in *T. timopheevii* has also been introduced into wheat and used in breeding. The gene *Pm6* derived from chromosome 2G and carried on chromosome 2B in wheat was reported in 1973 (Jorgensen and Jensen, 1973) and has proved durable to date (Wan et al., 2020). Resistance to diseases including spot blotch, tan spot, Stagonospora nodorum blotch, Septoria tritici blotch and loose smut has also been identified in *T. timopheevii* (Singh et al., 2006; Timonova et al., 2013) along with resistance to green bug (Rumyantsev et al., 2019) revealing it to be a rich source of resistance to a wide range of biotic stresses.

Resistance to FHB has been identified in a number of accessions of *T. timopheevii*. One of three accessions exhibited moderate type 2 resistance while all three lacked resistance to initial infection (type1) (Yong-Fang et al., 1997). FHB resistance derived from *T. timopheevii* has also been characterised in two separate studies. *Triticum timopheevii* accession PI 343447 was crossed and backcrossed to spring wheat Crocus and line TC 67 was selected on the basis of resistance to FHB and agronomic characteristics (Cao et al., 2009). TC 67 was crossed to the FHB susceptible variety Brio and a population of 230 F7 recombinant inbred lines produced and characterised for FHB resistance in glasshouse (type2) and field trials (incidence (type 1) and severity (type 2)) (Malihipour et al., 2017). Two QTL were identified on the long arm of chromosome 5A with one near the centromere in the interval between markers *cfd6*.*1* and *barc48* and the second more distal between *cfd39* and *cfa2185*. Both QTL contributed more than one FHB resistance trait. The QTL in the interval between *cfd6*.*1* and *barc48* was associated with reduced disease incidence and severity and reduced Fusarium damaged kernels (FDK) in field trials. The more potent QTL between *cfd39* and *cfa2185* was associated with reduced FDK in field trials and reduced severity in glasshouse trials following point inoculation indicating that it conferred type 2 resistance (Malihipour et al., 2017). Resistance to FHB derived from *T. timopheevii* was identified in a separate population developed between wheat line PI 277012 that contained *T. timopheevii* in its pedigree and wheat variety Grandin (PI 531005) (Chu et al., 2011). Two QTL contributing resistance to FHB in field trials and glasshouse point inoculation trials were identified on chromosome 5A with one on the short arm and one on the long arm. The QTL *Qfhb*.*rwg-5A*.*1* on the short arm resides in a similar location to *Qfhs*.*ifa-5A* identified in Sumai 3 with both being in the region of marker *XBarc180* (Chu et al., 2011). The QTL (*Qfhb*.*rwg-5A*.*2*) explaining most of the phenotypic variance flanks the QTL interval *cfd39* and *cfa2185* identified in TC 67 derived from *T. timopheevii* PI 343447 making it highly likely that these represent the same resistance (Chu et al., 2011). In both cases the QTL also flanked the *Q* locus with FHB resistance being associated with the *q* allele that prevents free-threshing reinforcing the view that the two QTL have a similar basis (Chu et al., 2011; Malihipour et al., 2016). It is unlikely that the differential FHB resistance is due to the *Q* locus itself because recombination between FHB resistance and non-free threshing was observed (Chu et al., 2011). The presence of the FHB resistance QTL on 5A indicates that they probably derive from chromosome 5A^t^ of *T. timopheevii*.

The FHB resistance of accession P95-99.1-1 appears to be predominantly of type 1 (resistance to initial infection) rather than type 2 (resistance to spread) as the greater resistance in this line compared to the other accessions tested was only evident following spray inoculation. This characteristic differentiates the resistance from that reported previously from *T. timopheevii* (Chu et al., 2011; Malihipour et al., 2016).

A large panel of interspecific hybrid lines has been developed of introgressions from *T. timopheevii* accession P95-99.1-1 in spring wheat Paragon. Advanced back-cross lines were further back-crossed with Paragon and self-fertilised to produce lines containing a relatively small number of introgressions (King et al., 2022). The number of introgressions retained within each line reduced with each back-cross with the exception of part of chromosome 2G (Devi et al., 2019). It has been demonstrated that chromosome 2G of *T. timopheevii* is preferentially transmitted accounting for its retention in lines over numerous back-crossings (Brown-Guedira et al., 1996). Assessment of FHB resistance requires relatively large numbers of plants and these need to be fixed for the presence of the introgression(s). Sufficient grain was available for only a proportion of the introgression panel and 32 of these carrying 57 unique introgressions from *T. timopheevii* (Figure 7) were assessed for resistance to FHB following spray inoculation.

Four introgression lines (Tim3, Tim4, Tim10 and Tim27) carry segments from 5A^t^ of *T. timopheevii* that are believed to cover the region associated with the potent FHB resistance QTL reported previously (Chu et al., 2011; Malihipour et al., 2017). In addition, the introgression carried by line Tim4 appears to cover the region containing the less potent resistance. None of these lines exhibited significantly greater FHB resistance than the wheat donor and it is concluded that accession P95-99.1-1 does not contain either of these FHB QTL.

Two lines (Tim5 and Tim6) were highly FHB resistant in the first year of testing. Both lines contain the preferentially transmitted segment of 2G and a segment of 3G equivalent to 3BS in wheat as revealed by KASP (Fig 7) and FISH (Fig 8) analysis. As many lines also contained the 2G segment but did not exhibit increased FHB resistance it was assumed that the resistance was conferred by the 3G segment. This was confirmed in the second year of trials. Line BC_2_F_4_-40 contains a 3G segment similar in size to that in Tim5 but lacks the preferentially transmitted segment of 2G. This line exhibited a similar high level of FHB resistance to Tim5 in the second year of trials. The size of the 3G segment in Tim5 and line BC_2_F_4_-40 is much smaller than that in Tim6, 48.8 Mb and 762.2 Mb respectively. Tim2, Tim24 and Tim35 all contain small segments (up to 10.75 Mb) of 3G introgressed onto the distal end of 3B but none of these lines showed enhanced FHB resistance in either trial (Figure 5, Figure 6). It is assumed, therefore, that the region associated with FHB resistance on the distal portion of 3BS is contributed by the equivalent region of 3G in the interstitial 38.05 Mb region between 10.75 and 48.8 Mb.

**Figure 6.**
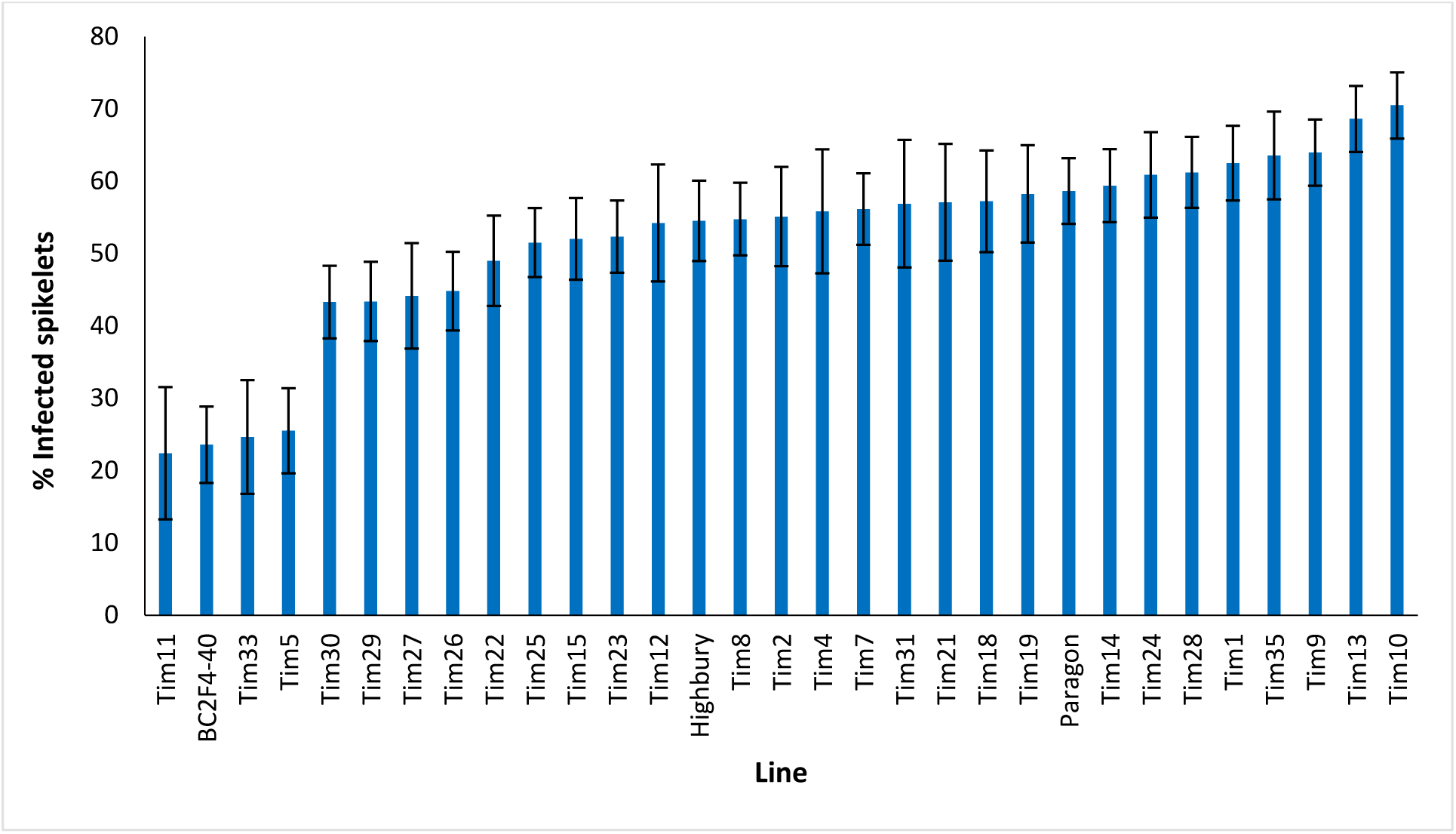
Visual FHB disease score from 2021 trial of 29 *T. timopheevii* accession P95-99.1-1/wheat introgression lines, Highbury and Paragon susceptible controls expressed as percentage of infected spikelets 21 days post spray inoculation with *F. culmorum* (1×10^5^ conidia per ml).

**Figure 7.**
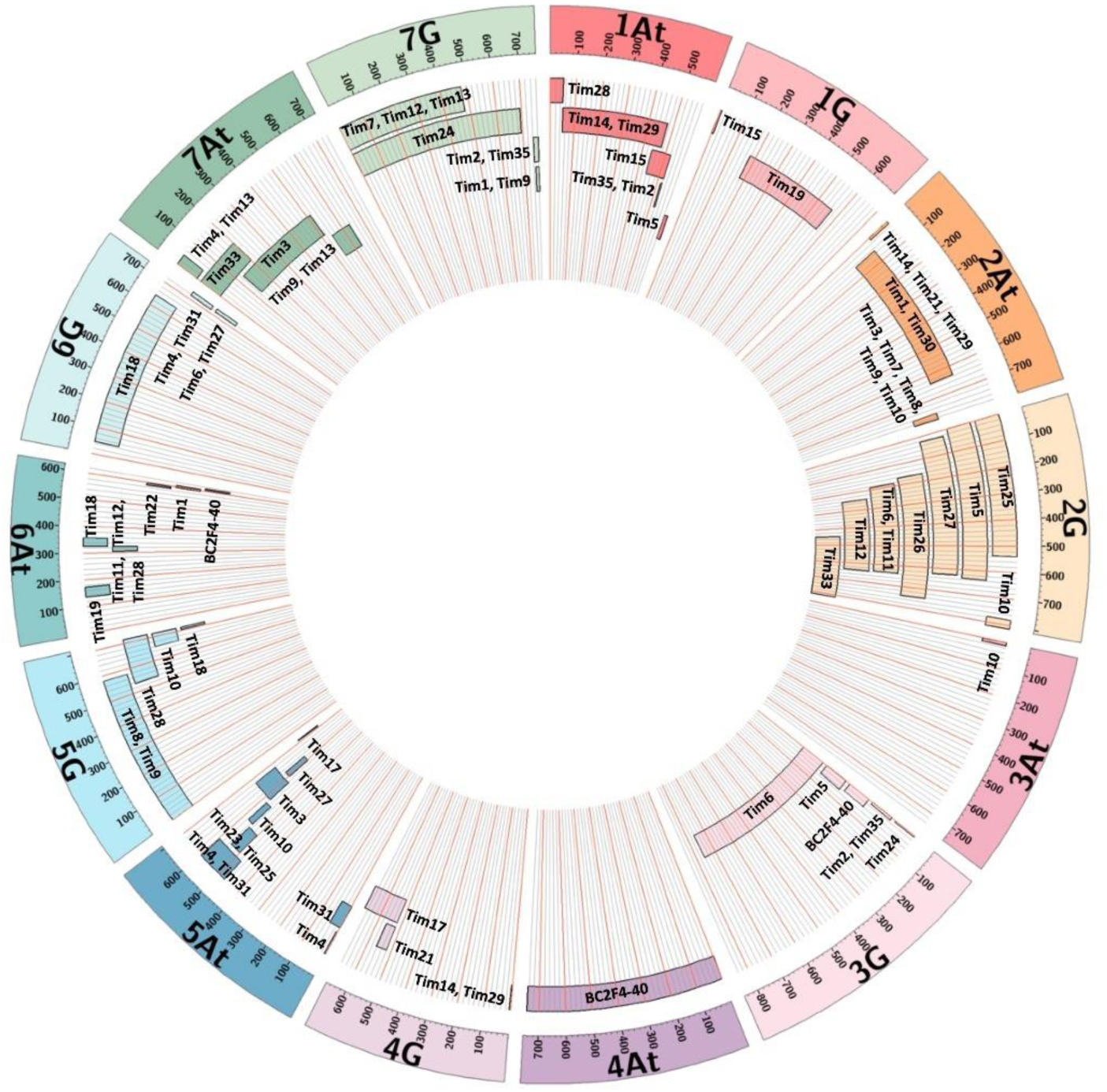
Representation of *T. timopheevii* chromosome segments contained within each Tim introgression line showing approximate position and size of each segment relative to each *T. timopheevii* chromosome.

**Figure 8.**
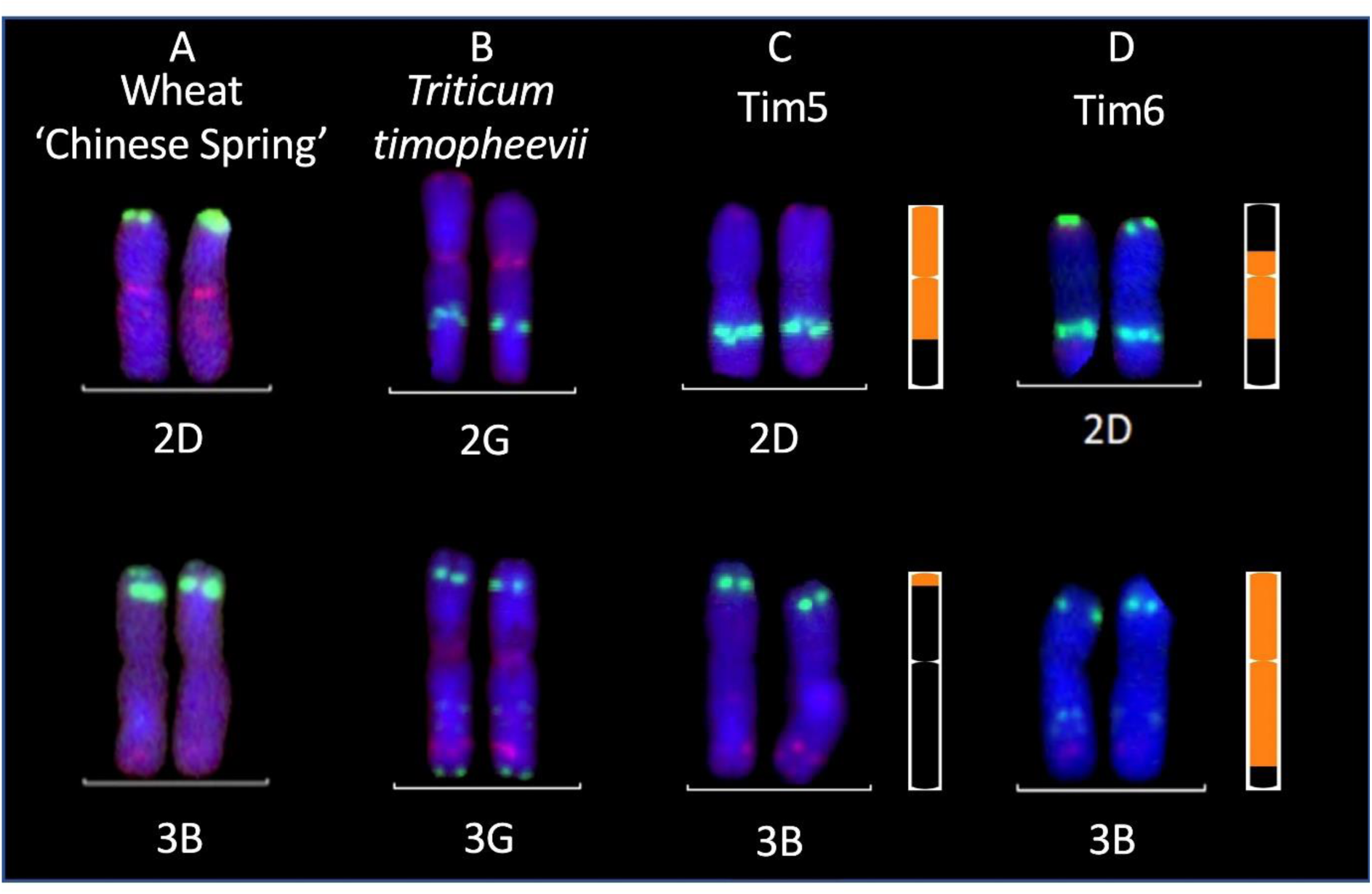
Multi-colour fluorescence *in situ* hybridisation analysis of root metaphase spreads of 2D and 3B chromosomes from (A) wheat cv. Chinese Spring (B) chromosomes 2G and 3G of *T. timopheevii* and chromosomes 2D and 3B of FHB resistant wheat-*T. timopheevii* introgression lines (C) Tim 5 and (D) Tim6. Green and red signals show pSc119.2 and pAs.1 binding sites, respectively. KASP marker-derived ideograms of the introgressions in Tim 5 and Tim 6 are shown to the right. Size of introgressions from A^t^ and G subgenomes is indicated in orange in a wheat chromosome shown in black.

Line Tim2 used in the FHB trial in 2020 contained segments of chromosomes 1A^t^ and 7G from *T. timopheevii* and segregated for loss of the short arm of chromosome 3B (3BS). The loss of 3BS was fixed in this line in the FHB trial in 2021. It has been postulated that the resistance conferred by *Fhb1* on chromosome 3BS is due to a loss of function (Su et al., 2019) of the histidine rich calcium binding protein. No significant increase in FHB resistance in Tim2 was observed in either year indicating that loss of 3BS does not result in increased FHB resistance. This observation is in agreement with the finding that loss of 3BS (Ma et al., 2006) or replacement of 3BS with 3HS (Hales et al., 2020) does not result in an increase in FHB resistance.

Two other lines showed enhanced FHB resistance in the 2021 trial. Tim11 was moderately resistant in the first year of trials and was one of the most resistant in the second trial when disease pressure was higher (Figs 5 and 6). This line contains segments from 2G and 6A^t^. The 6A segment is similar to that in a number of other lines and the 2G segment is present in Tim5. None of the lines containing either the 2G and 6A^t^ segments showed enhanced resistance and so the origin of this resistance is unclear. It is possible that this line contains additional segments of *T. timopheevii* chromosome that are too small to detect with either the SNP markers or with FISH. This is particularly relevant for introgressions from the A^t^ genome. Recombination between the A genome of wheat and the A^t^ genome of *T. timopheevii* is much more prevalent than that between the B or D genome of wheat and the G genome of *T. timopheevii* (Timonova et al., 2013). The size of A^t^ introgressions may be reduced during the process of backcrossing to reduce and stabilise the number of introgressions and so their presence may not be detected using the current SNP marker set.

Insufficient grain of Tim 33 was available for FHB resistance assessment in 2020 but this line exhibited a high level of resistance in the 2021 trial (Fig. 6). Tim33 contains the preferentially transmitted segment of 2G and a segment on 7AS presumed to originate from 7A^t^ of *T. timopheevii*. The segment on 7AS of Tim33 is in the region between 0 to ∼200Mb. Two other lines (Tim4 and Tim13) also possess segments introduced to 7AS but the size of the introgression (0- ∼42Mb) is considerably less than that in Tim33. In addition, Tim3 contains a segment introduced in the region 128-515Mb on 7A. None of these lines exhibited high levels of FHB resistance and so it is concluded that the resistance exhibited by Tim33 is due to genes present in the region between ∼42 and 128 Mb. Additional testing of these lines is required to confirm the presence and location of the FHB resistance conferred by this interval of 7A^t^.

*Triticum timopheevii* is a rich source of diversity for the introduction of beneficial traits from the secondary gene pool of wheat. We have identified an accession of *T. timopheevii* with very high levels of type 1 FHB resistance. The resistance appears to be novel and is expressed when introduced into wheat. Introgression of individual chromosomal segments (e.g 38.05 Mb region between 10.75 and 48.8 Mb of 3G) significantly increase FHB resistance in wheat. The material generated within this study provides a new source of FHB resistance for evaluation in wheat breeding programmes.

## Conflict of Interest

The authors declare that the research was conducted in the absence of any commercial or financial relationships that could be construed as a potential conflict of interest.

## Author contribution

IK, JK and PN conceived this work. JK, CY and SG developed the introgression lines reported here. SG carried out the genotyping and CY carried out the FISH analysis of introgression lines. AS, MT and PN carried out the disease screening of plants and AS analysed the data produced. PN, AS, IK, JK, CY and SG contributed to writing this manuscript.

## Funding

This work was supported by the UK Biotechnology and Biological Sciences Research Council (BBSRC) through the Designing Future Wheat [BB/P016855/1] Institute Strategic Programme.

## Acknowledgements

AS and PN would like to thank the Horticultural Services team of the John Innes Centre for their input into care of the plants used in these experiments.

## Notes

### Competing Interest Statement

The authors have declared no competing interest.

